# ODNA: Identification of Organellar DNA by Machine Learning

**DOI:** 10.1101/2023.01.10.523051

**Authors:** Roman Martin, Minh Kien Nguyen, Nick Lowack, Dominik Heider

## Abstract

**Motivation:** Identifying organellar DNA, such as mitochondrial or plastid sequences, inside a whole genome assembly, remains challenging and requires biological background knowledge. To address this, we developed ODNA based on genome annotation and machine learning to fulfill.

**Results:** ODNA is a software that performs organellar DNA sequence classification of a genome assembly by machine learning based on a pre-defined genome annotation workflow. We trained our model with 829,769 DNA sequences from 405 genome assemblies and achieved very high predictive performance (e.g., MCC of 0.61) on independent validation data, thus outperforming existing approaches significantly.

**Availability:** Our software ODNA is freely accessible as a web service at https://odna.mathematik.uni-marburg.de and can also be run in a docker container. The source code can be found at https://gitlab.com/mosga/odna and the processed data at Zenodo (DOI: 10.5281/zenodo.7506483).

## Introduction

A comprehensive genome of a eukaryotic species includes organellar DNA such as mitochondria, chloroplastid, or even some other plastid’s DNA. Except for some rare cases (1), most eukaryotic cells contain at least a mitogenome. In practice, organellar DNA is often sequenced together with chromosomal genome sequencing approaches, as visualized in **Figure S1**. Based on the selected software, highly abundant organelles sequences can affect genome analyses. Therefore, methods were already developed to physically reduce putative contamination by organelles (2). However, identifying organellar DNA by computational methods can also help to reduce the problem. Additionally, the preservation and identification of organellar DNA sequences can be further beneficial for performing phylogenetics or taxonomy inference analyses (3, 4). However, the identification remains challenging, especially if the samples are highly contaminated with bacterial sequences that can contain similar genes to those from organelles.

In most cases, there is no single rule to precisely determine if an organellar DNA sequence is within a genome assembly. Through evolution, for example, the mitogenome differentiates massively through different taxonomic clades. The mitogenome can be compact, with 11 kilobases (kb) in some animals, while reaching up to 1.1 Mb in some plants (5). Even analyzing the gene composition is not sufficient enough to distinguish the correct sequences, since genes have transferred through millions of years (6), ending up in a set of highly conserved genes. In recent years, several tools have been developed that specifically identify organellar DNA, such as the MitoFinder (7) for mitochondrial DNA or the chloroExtractor (8) for chloroplasts. Generally, using raw sequencing reads instead of a genome assemblies is recommended, but not always provided. While most organelles tools use raw sequencing data, the MitoFinder can use both genomes assembly and raw sequencing reads. Furthermore, it was recently shown that it is possible to use genome annotation data from MOSGA 2 (9) based on a combination of different multiple prediction tool outputs to identify some organellar DNA.

Here, we present the software ODNA and evaluate its performance. Using machine learning (ML), ODNA classifies sequences from a given eukaryotic genome assembly into nucleolar or organellar DNA. Technically, the software is a pipeline based on a pre-defined Snakemake (10) workflow and MOSGA (9, 11) that embeds an additional ML model for the classification.

## Materials and Methods

We performed hundreds of eukaryotic genome annotations and noted which sequences belong to organelles according to the NCBI organelles database. We completed ML training based on these annotations to obtain an accurate model. The data retrieval procedure is visualized in **Figure S2**. As a result, we developed ODNA, a minimalized pre-defined genome annotation software based on MOSGA, which gathers the same annotation features and includes the best ML model. ODNA can classify if a sequence inside a genome assembly belongs to organellar origin.

### Machine Learning

As the feature set, ODNA annotates for each sequence in each genome assembly the repeating elements via Red (12), the ribosomal RNAs with barrnap, transfer RNAs with tRNAScan-SE 2 (13), CpG islands with newcpgreport from the EMBOSS suite (14), and DIAMOND (15) searches against a mitochondrial and plastid gene databases from MOSGA 2 (9). Additionally, characteristics such as the GC content, sequence length, and substantial deviation from the average GC content in an assembly were encoded, as well as the density of the most features per 1 Mb. We used a stratified training-to-test ratio of 1:5 for the 10-fold cross-validation (CV) with various ML models provided by scikit-learn, including Linear Classifiers, Random Forests, k-nearest neighbors, and AdaBoost. We evaluated our model on an independent validation dataset consisting of 14,514 sequences from ten eukaryotic genome assemblies in a real-world use-case, and compared the results to Mitofinder. A more detailed description is provided in the supplementary information. All data and scripts are available on Zenodo, ensuring reproducibility.

### Comparison

According to our knowledge, no similar software like ODNA is freely available. Therefore, we compared the classification performance of ODNA with MitoFinder, since both software can use eukaryotic genome assemblies to identify mitochondrial sequences. For the comparison, we used a validation set of 14,514 sequences from ten eukaryotic genome assemblies that were not included in the CV (see **Table S1**).

## Results

In total, 405 eukaryotic genomes with 829,769 sequences were annotated for training the ML model. Among these sequences, 450 are organellar sequences. The best ML model (AdaBoost) has an MCC of 0.90 for our test data (**Figure S3**). Although we did not include any sequences in the training or test data from the taxonomical class *Bigyra*, our generalized model was capable of identifying the mitochondrial sequence, for example, from *Cafeteria burkhardae* (16).

A comparison between MitoFinder and ODNA on our validation data reveals that ODNA outperforms MitoFinder with an MCC of 0.61 versus 0.37, mainly deriving from more true positives and fewer false positives as shown in Tab. 1. Additionally, the median execution time was shorter, with 24 min instead of 141 min.

**Table 1.** Performance comparison between MitoFinder and ODNA. The true positives (TP), true negatives (TN), false positives (FP), false negatives (FN), Matthews correlation coefficient (MCC), F1-score (F1), precision (Pre), sensitivity (Sen), specificity (Spe) and execution time are compared.

| Software | TP | TN | FP | FN | MCC | F1 | Pre | Sen | Spe | $\tilde{t}$ (s) |
| --- | --- | --- | --- | --- | --- | --- | --- | --- | --- | --- |
| MitoFinder | 7 | 14,476 | 28 | 3 | 0.37 | 0.31 | 0.20 | 0.70 | 1.00 | 8,446 |
| ODNA | 10 | 14,487 | 17 | 0 | 0.61 | 0.54 | 0.37 | 1.00 | 1.00 | 1.447 |

## Discussion

With ODNA, we provide an easy-to-use, web-interface tool that does not require any parametrization and identifies organellar DNA precisely across all eukaryotic clades via ML. Compared to MitoFinder, ODNA is faster and more precise, demonstrating how meta-information from a DNA sequence can be used to identify meaningful biological properties. However, for the sake of completeness, we want to mention that MitoFinder’s main focus lies in identifying mitochondrial sequences and their annotation. Furthermore, since the technical pipeline is established, we consider updating our ODNA model if the NCBI genome assembly database grow.

## ACKNOWLEDGEMENTS

This work was supported by the BMBF-funded de.NBI Cloud within the German Network for Bioinformatics Infrastructure (de.NBI).

