## Supplementary Information for "ODNA: Identification of Organellar DNA by Machine Learning"

#### This PDF file includes:

Supplementary text
Tables S1 to S4
Figs. S1 to S10

### Supplementary Information Text

#### Data Retrieval, Encoding and Data Analysis

The data used for the machine learning approach was retrieved from NCBI. First, we downloaded all data from the NCBI Organellar Sequences Database, release number 214 (see Figure S2). Second, we downloaded all genome assemblies from NCBI Genbank, referenced through the Organellar Sequence Database. Then, we generated the machine learning data by performing 405 genome annotations via MOSGA 2 (see Figure S4), resulting in 829,769 annotated sequences. Subsequently, we developed ODNA (see Figure S5) as a modified fork from MOSGA v.2.1.5 with an integrated machine learning model.

To encode the data, we used multiple tools, filtered the results (see ODNA parameter), and computed the features listed in Table S3. A Spearman and Pearson correlation analysis did not reveal a clear correlation between the target variable **Organelle** and any feature (see Figure S6), indicating that more than one feature is required to classify the organellar DNA sequence.

Although the Random Forest model was well-performing but in the end not used for the ODNA since AdaBoost performed better, the visualization of the mean decrease in impurity of the Random Forest model in Figure S7 from the Random Forest classifier shows convincing results regarding the feature's contribution. The features Mito, Plastid, their density MD and PD have the greatest effect. To analyze the influence of these features on the the model's performance, we removed the features Mito, Plastid, MD, and PD in one approach, resulting in worse classification performance. We could observe a similar reduced performance using only these features. Only the combination of all features resulted in the best-performing models. Interestingly, MitoFinder uses primarily a database of mitochondrial genes to identify the mitogenome, which is similar to our **Mito** feature.

Furthermore, we analyzed the taxonomical association of each genome assembly used for machine learning (see Figure S8). Less surprisingly, we can confirm that there is an overrepresentation of available genome data for some taxas, such as Chordata. This imbalance in the availability of genomic data is already well-described in the literature.

#### Machine Learning

The scikit-learn library carried out all machine learning computations within a Jupyter-Notebook in Python 3. The data was split in a stratified fashion, resulting in 663,455 training (80 %) and 165,864 test (20 %) data entries. A standard scaler for all data was fitted, and training was performed using 10-fold cross-validation. To evaluate the models' performance, we used trained for the best Matthews correlation coefficient (MCC) according to Equation 1.

$$\text{MCC} = \frac{TP \cdot TN - FP \cdot FN}{\sqrt{(TP + FP) \cdot (TP + FN) \cdot (TN + FP) \cdot (TN + FN))}} \quad [1]$$

The hyper-parameterization was performed within the cross-validation (see Figure S9) and realized through a Halving Grid Search and 100 Randomized Searches. To be unbiased to specific kinds of machine learning, we tested multiple machine learning models, including AdaBoost, Bagging, Decision Tree, Extra Trees, Gradient Boosting, K-nearest neighbor, Multi-layer perceptron (MLP), and Stochastic Gradient Descent (SGD).

The best model was an AdaBoost Classifier derived from the Halving Grid Search hyper-parameterization. It has an MCC of 0.90 and an F1-Score of 0.90 (see Figure S3) with the test dataset. The best model was an AdaBoost Classifier derived from the Halving Grid Search hyper-parameterization. It has an MCC of 0.90 and an F1-Score of 0.90 (see Figure S3) with the test dataset. We also tested our model in a real-world use case with ten eukaryotic genome assemblies across all different taxonomical clades with 14,514 sequences, which we considered the validation dataset.

Additionally, to ensure that our model does not apply only to the sequences from already learned taxa, we tested and successfully classified the mitochondrial DNA from the protist Cafeteria roenbergensis (GCA_008330645.1), which the whole taxonomical class no data were learned.

#### Benchmark

In order to compare the capability of ODNA and MitoFinder in identifying organellar DNA sequences, we randomly selected ten genome assemblies that were used neither in training nor during machine learning (see Machine Learning section). Since we assume that both tools would be able to identify the organelles or, respectively, the mitochondrial DNA, we tried to minimalize the parameterization. Sequences within a genome assembly were ignored for the evaluation if they belong to other organelles except for mitochondria. All used software is listed in Table S4.

**MitoFinder parameter.** MitoFinder requires the preparation of a reference gene database. We generated a new reference database without a taxonomic restriction, as recommended by the developers.

```
esearch -db nuccore -query "\"mitochondrion\"[All Fields] AND (refseq[filter] AND mitochondrion[filter]
AND (\"12000\"[SLEN] : \"20000\"[SLEN]))" | efetch -format gbwithparts > reference.gb
```

We run this command exemplary for each genome assembly. Here for GCF_000001215.4.

```
~/MitoFinder/mitofinder -j GCF_000001215.4 -a ~/VALIDATION/genomes/GCF_000001215.4.fasta -r
~/VALIDATION/reference.gb -p 32 -o 1 --ignore --override
```

**ODNA parameter.** The beauty of ODNA lies in the fact that it is neither necessary nor possible to parameterize ODNA. It is pre-configured, and user can only provide a valid genome FASTA or a genebank flat (GBFF) file with sequences. However, the underlying tools are parameterized in the sense as MOSGA 2 allows parameterization. Red: Repeats were only considered with the minimal length of 100 bases. tRNAScan-SE 2.0 transfer-RNAs with a minimal score of 70. barrnap: only rRNAs from the eukaryota kingdom, length cut off of 80, rejection rate of 25 and a minimal evalue of $10^{-6}$.

### Computational resource

We used two servers from the German governmental deNBI cluster to perform all MOSGA genome annotations, each with 32 CPU cores and 64 GB of main memory. The required genome annotation time for the dataset creation is plotted against the genome size in Figure S10. Additionally, the machine learning computations, and the MitoFinder and ODNA executions were run independently on one of these machines.

**Table S1. Comparison of the prediction performance of MitoFinder and ODNA with different genome assemblies. A total of 14,514 sequences were tested from ten different genome assemblies. The genome assembly, species, number of sequences, true positives (TP), true negatives (TN), false positives (FP), false negatives (FN), and execution time are listed below.**

| | | | MitoFinder | | | | | ODNA | | | | |
|---|---|---|---|---|---|---|---|---|---|---|---|---|
| Assembly | Species | Sequences | TP | TN | FP | FN | Time (s) | TP | TN | FP | FN | Time (s) |
| GCF_000001215.4 | *Drosophila melanogaster* | 1,870 | 1 | 1,860 | 9 | 0 | 8,757 | 1 | 1,869 | 0 | 0 | 279 |
| GCF_000002335.3 | *Tribolium castaneum* | 2,082 | 1 | 2,074 | 7 | 0 | 7,299 | 1 | 2,076 | 5 | 0 | 359 |
| GCF_001856785.1 | *Myzus persicae* | 4,021 | 1 | 4,013 | 7 | 0 | 9,131 | 1 | 4,013 | 7 | 0 | 613 |
| GCF_002099425.1 | *Phascolarctos cinereus* | 1,970 | 1 | 1,965 | 4 | 0 | 10,849 | 1 | 1,966 | 3 | 0 | 10,397 |
| GCF_002880755.1 | *Pan troglodytes* | 4,346 | 1 | 4,344 | 1 | 0 | 10,446 | 1 | 4,343 | 2 | 0 | 28,272 |
| GCA_002994745.2 | *Rosa chinensis* | 54 | 0 | 53 | 0 | 1 | 8,175 | 1 | 53 | 0 | 0 | 2,280 |
| GCF_003290485.1 | *Malassezia restricta* | 10 | 0 | 9 | 0 | 1 | 7,584 | 1 | 9 | 0 | 0 | 39 |
| GCF_015533775.1 | *Penicillium roqueforti* | 84 | 0 | 83 | 0 | 1 | 7,415 | 1 | 83 | 0 | 0 | 79 |
| GCF_019923935.1 | *Bubalus bubalis* | 26 | 1 | 25 | 0 | 0 | 8,881 | 1 | 25 | 0 | 0 | 41,169 |
| GCF_900324465.2 | *Anabas testudineus* | 51 | 1 | 50 | 0 | 0 | 7,872 | 1 | 50 | 0 | 0 | 2,360 |
| | | $\Sigma$ | 7 | 14,476 | 28 | 3 | | 10 | 14,487 | 17 | 0 | |
| | | $\widetilde{t}$ | | | | | **8,466** | | | | | **1,447** |

**Table S2. List of all annotated genome assemblies used for the machine learning.**

| Genome assemblies | | | | | | |
|---|---|---|---|---|---|---|
| GCF_002072235.1 | GCF_017589495.1 | GCF_020740795.1 | GCF_005870125.1 | GCF_020615455.1 | GCF_006386435.1 | GCF_000230625.1 |
| GCF_000150675.1 | GCF_018340385.1 | GCF_020360975.1 | GCF_005239225.1 | GCF_000004255.2 | GCF_001901225.1 | GCF_012393455.1 |
| GCF_021130785.1 | GCF_018687715.1 | GCF_000328475.1 | GCF_018350155.1 | GCF_000347755.3 | GCF_008403515.1 | GCF_001417885.1 |
| GCF_002114115.1 | GCF_001298625.1 | GCF_017976325.1 | GCF_008692025.1 | GCF_000764305.2 | GCF_010645085.1 | GCF_000002415.2 |
| GCF_000972845.2 | GCF_000003025.6 | GCF_003987935.1 | GCF_014108415.1 | GCF_009762305.2 | GCF_019740435.1 | GCF_023101765.2 |
| GCF_011064425.1 | GCF_004329235.1 | GCF_009730915.1 | GCF_000182895.1 | GCF_000504015.1 | GCF_900324485.2 | GCF_016881025.1 |
| GCF_002021735.2 | GCF_001659605.2 | GCF_000313985.2 | GCF_003576645.1 | GCF_022985175.1 | GCF_020536065.1 | GCF_009819535.1 |
| GCF_000182925.2 | GCF_000150975.2 | GCF_900185275.1 | GCF_000003225.4 | GCF_000003525.1 | GCF_019186545.1 | GCF_003073045.1 |
| GCF_013396075.1 | GCF_905237075.1 | GCF_017639785.1 | GCF_014843425.1 | GCF_001723895.1 | GCF_018812025.1 | GCF_000741045.1 |
| GCF_905475465.1 | GCF_004916995.1 | GCF_900002335.2 | GCF_003339765.1 | GCF_002078875.1 | GCF_900002385.2 | GCF_022113875.1 |
| GCF_007399415.2 | GCF_000280675.1 | GCF_023721935.1 | GCF_004153795.1 | GCF_002220235.1 | GCF_014754425.2 | GCF_019202705.1 |
| GCF_015227675.2 | GCF_000149985.1 | GCF_014066315.1 | GCF_000611645.1 | GCF_000188115.5 | GCF_022379125.1 | GCF_024362695.1 |
| GCF_004011695.1 | GCF_017639515.1 | GCF_922984935.1 | GCF_000956335.1 | GCF_001625215.1 | GCF_014570555.1 | GCF_007474595.2 |
| GCF_900964775.1 | GCF_014633375.1 | GCF_000214015.2 | GCF_013052645.1 | GCF_002201575.1 | GCF_023701775.1 | GCF_000388065.1 |
| GCF_004193775.1 | GCF_015220715.1 | GCF_017639745.1 | GCF_900634795.3 | GCF_023343755.1 | GCF_905220565.1 | GCF_900090045.1 |
| GCF_000149555.1 | GCF_003918875.1 | GCF_018492685.1 | GCF_008315115.2 | GCF_006542625.1 | GCF_000648675.2 | GCF_014898055.1 |
| GCF_900681995.1 | GCF_000149925.1 | GCF_013358625.1 | GCF_016086655.3 | GCF_017311325.1 | GCF_020740605.2 | GCF_004115215.2 |
| GCF_019059575.1 | GCF_017639655.2 | GCF_018350215.1 | GCF_900005765.1 | GCF_007655135.1 | GCF_000149035.1 | GCF_022539355.2 |
| GCF_013396195.1 | GCF_000341935.2 | GCF_000143365.1 | GCF_024586455.1 | GCF_001853355.1 | GCF_014083535.2 | GCF_902167405.1 |
| GCF_001949145.1 | GCF_000238955.4 | GCF_016746365.2 | GCF_000184455.2 | GCF_014466165.1 | GCF_002571385.1 | GCF_020975775.1 |
| GCF_002251995.1 | GCF_900747795.1 | GCF_022830035.1 | GCF_019202785.1 | GCF_002922805.2 | GCF_002443255.1 | GCF_018296145.1 |
| GCF_006345805.1 | GCF_009819705.1 | GCF_023278565.1 | GCF_003713225.1 | GCF_000231095.2 | GCF_003789085.1 | GCF_018345385.1 |
| GCF_016077235.1 | GCF_021613505.1 | GCF_003709585.1 | GCF_004382145.1 | GCF_000149205.2 | GCF_000281125.3 | GCF_003957565.2 |
| GCF_002776465.1 | GCF_000004075.3 | GCF_004382195.2 | GCF_000149685.1 | GCF_000004695.1 | GCF_012559485.2 | GCF_900005855.1 |
| GCF_014905175.1 | GCF_011762595.1 | GCF_011075155.1 | GCF_022539315.1 | GCF_002938485.1 | GCF_000165395.1 | GCF_002263795.2 |
| GCF_910594005.1 | GCF_011004845.1 | GCF_902362185.1 | GCF_018350195.1 | GCF_014851395.1 | GCF_006381635.1 | GCF_000293215.1 |
| GCF_002926055.2 | GCF_018977255.1 | GCF_900700415.2 | GCF_003640425.2 | GCF_020631705.1 | GCF_002910315.2 | GCF_017654675.1 |
| GCF_000002285.5 | GCF_015220235.1 | GCF_004027225.2 | GCF_016859285.1 | GCF_011764305.1 | GCF_009663435.1 | GCF_004353265.1 |
| GCF_002775205.1 | GCF_001406875.1 | GCF_003254395.2 | GCF_900626175.1 | GCF_001640805.2 | GCF_008121235.1 | GCF_000149335.2 |
| GCF_003704095.1 | GCF_001879475.1 | GCF_005190385.1 | GCF_902500615.1 | GCF_003597395.1 | GCF_017312705.1 | GCF_014066325.1 |
| GCF_001577835.2 | GCF_004193835.1 | GCF_019578655.1 | GCF_015832195.1 | GCF_905147795.1 | GCF_010993605.1 | GCF_900002375.2 |
| GCF_005281545.1 | GCF_001039355.2 | GCF_013100865.1 | GCF_003009895.1 | GCF_014805685.1 | GCF_002127325.2 | GCF_000145635.1 |
| GCF_902655055.1 | GCF_003327715.1 | GCF_000150115.1 | GCF_932294415.1 | GCF_000499545.2 | GCF_013377495.2 | GCF_009870125.1 |
| GCF_016617805.1 | GCF_001442555.1 | GCF_001854935.1 | GCF_000146045.2 | GCF_001890085.1 | GCF_011800145.1 | GCF_009650485.1 |
| GCF_019176455.1 | GCF_000317415.1 | GCF_008632895.1 | GCF_013122585.1 | GCF_014805655.1 | GCF_007990345.1 | GCF_008831285.2 |
| GCF_000002545.3 | GCF_008728515.1 | GCF_010389155.1 | GCF_000150825.1 | GCF_020184175.1 | GCF_000143415.4 | GCF_015586225.1 |
| GCF_912470025.1 | GCF_022581195.2 | GCF_002234675.1 | GCF_943737965.1 | GCF_902201215.1 | GCF_004324835.1 | GCF_016699345.2 |
| GCF_000151145.1 | GCF_000003745.2 | GCF_018924745.1 | GCF_000350225.1 | GCF_016077325.2 | GCF_023343835.1 | GCF_023158985.2 |
| GCF_000002775.4 | GCF_013347855.1 | GCF_001649575.2 | GCF_003368295.1 | GCF_020740685.1 | GCF_000002235.5 | GCF_014356525.1 |
| GCF_016861625.1 | GCF_017499595.1 | GCF_009764315.1 | GCF_004664715.2 | GCF_000002985.6 | GCF_009873245.2 | GCF_017591425.1 |
| GCF_008641045.1 | GCF_018143015.1 | GCF_016746245.2 | GCF_900497805.2 | GCF_013402915.1 | GCF_004354835.1 | GCF_014108235.1 |
| GCF_002157705.1 | GCF_009829125.1 | GCF_019609905.1 | GCF_002880775.1 | GCF_003724035.1 | GCF_020379485.1 | GCF_021134715.1 |
| GCF_001654055.1 | GCF_014570535.1 | GCF_024364675.1 | GCF_000182765.1 | GCF_000346465.2 | GCF_020497125.1 | GCF_000002865.3 |
| GCF_001540865.1 | GCF_022316705.1 | GCF_007565055.1 | GCF_016545825.1 | GCF_900246225.1 | GCF_002079055.1 | GCF_000149615.1 |
| GCF_000165365.1 | GCF_016509475.1 | GCF_000149245.1 | GCF_000303195.2 | GCF_009769625.2 | GCF_003342905.1 | GCF_016745375.1 |
| GCF_014805675.1 | GCF_000146945.2 | GCF_003331165.1 | GCF_015476345.1 | GCF_000006445.2 | GCF_015237465.2 | GCF_020740725.1 |
| GCF_020171115.1 | GCF_023653815.1 | GCF_002917755.1 | GCF_900963305.1 | GCF_000836215.1 | GCF_023373825.1 | GCF_023119035.1 |
| GCF_901001165.1 | GCF_000004515.6 | GCF_016699485.2 | GCF_009389715.1 | GCF_000699065.1 | GCF_003086295.2 | GCF_000208745.1 |
| GCF_000687475.1 | GCF_023065955.1 | GCF_018320785.1 | GCF_016772045.1 | GCF_017654505.1 | GCF_013373865.1 | GCF_009805555.1 |
| GCF_010909765.2 | GCF_017591435.1 | GCF_011397635.1 | GCF_000091025.4 | GCF_016746395.1 | GCF_016432855.1 | GCF_022458985.1 |
| GCF_003255815.1 | GCF_009819795.1 | GCF_008122165.1 | GCF_907164915.1 | GCF_000146915.1 | GCF_004118075.2 | GCF_013265735.2 |
| GCF_905116875.1 | GCF_000003855.2 | GCF_000002425.4 | GCF_000237925.1 | GCF_014898765.1 | GCF_000240135.2 | GCF_000239435.1 |
| GCF_000003815.2 | GCF_000342415.1 | GCF_000090985.2 | GCF_000151545.1 | GCF_000003515.1 | GCF_937001465.1 | GCF_010015445.1 |
| GCF_016920845.1 | GCF_004379255.1 | GCF_016801865.1 | GCF_009834535.1 | GCF_017590055.1 | GCF_016835505.1 | GCF_008692095.1 |
| GCF_020382885.2 | GCF_900880675.1 | GCF_002926085.2 | GCF_020085105.1 | GCF_014839805.1 | GCF_910596095.1 | GCF_003343065.1 |
| GCF_000149585.1 | GCF_018831695.1 | GCF_019320065.1 | GCF_002706865.2 | GCF_901000725.2 | GCF_021292245.1 | GCF_002863925.1 |
| GCF_003121395.1 | GCF_902148845.1 | GCF_001186385.1 | GCF_001858045.2 | GCF_916048215.2 | GCF_000309985.2 | GCF_000006355.2 |
| GCF_001444195.1 | GCF_000151425.1 | GCF_011125445.2 | GCF_002042975.1 | GCF_013358895.1 | GCF_000004195.4 | |

**Table S3. List of all features used. The feature name, data type, software source, and comprehensive meaning are listed in the table below.**

| Features | Data type | Source | Meaning |
|---|---|---|---|
| Repeats | integer | Red | number of repeats |
| tRNA | integer | tRNAScan-SE | number of transfer RNAs |
| rRNA | integer | barrnap | number of ribosomal RNAs |
| rRNAp | integer | barrnap | number of partial ribosomal RNAs |
| Mito | integer | MOSGA 2 | number of matches against our mitochondrial gene database |
| Plastid | integer | MOSGA 2 | number of matches against our plastid gene database |
| CpGisland | integer | newcpgreport | number of CpG islands |
| GC | float | MOSGA 2 | percental GC content |
| GC-dev | float | MOSGA 2 | percental GC content deviation from the assembly average |
| GC-out | boolean | MOSGA 2 | True if GC content is higher or lower than the averaged GC content of an assembly |
| Len | float | MOSGA 2 | length in kilobases (kb) |
| MD | float | MOSGA 2 | mitochondrial gene matches (Mito) per 10 kb |
| PD | float | MOSGA 2 | plastid gene matches (Plastid) per 10 kb |
| RD | float | ODNA | Repeats per 10 kb |
| TD | float | ODNA | tRNAs per 10 kb |
| RRD | float | ODNA | rRNAs per 10 kb |
| CpGD | float | ODNA | CpG islands per 10 kb |
| Fraction | float | ODNA | (sequence length / total assembly length) * 1000 |
| Organelle | boolean | NCBI | target variable |

**Table S4. List of the used software and versions.**

| Software | Version |
|---|---|
| MOSGA | v.2.1.5 |
| Red | 05/22/2015 |
| newcpgreport | EMBOSS:6.6.0.0 |
| tRNAscan-SE | 2.0.8 |
| barrnap | 0.9 |
| scikit-learn | 1.1.3 |
| jbrowse | 1.5.1 |
| tbl2asn | 25.8 |

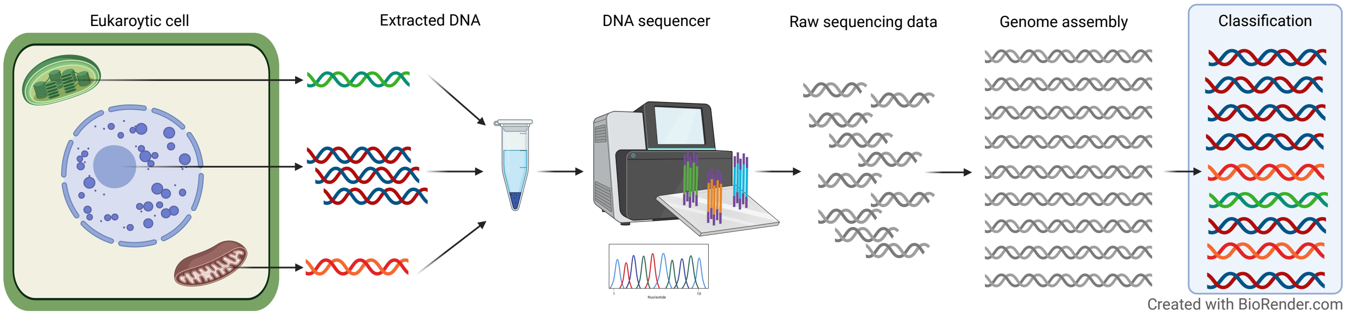


**Fig. S1. Visualization of the technical workflow**. Based on the selected DNA extraction method, organellar DNA will be extracted for a sequencing procedure and require special treatment during the annotation and submission. However, depending on the data quality and reference knowledge, it is challenging to differentiate which sequence derives from an organelle. ODNA presents software that can classify organellar DNA based on specific annotation patterns.

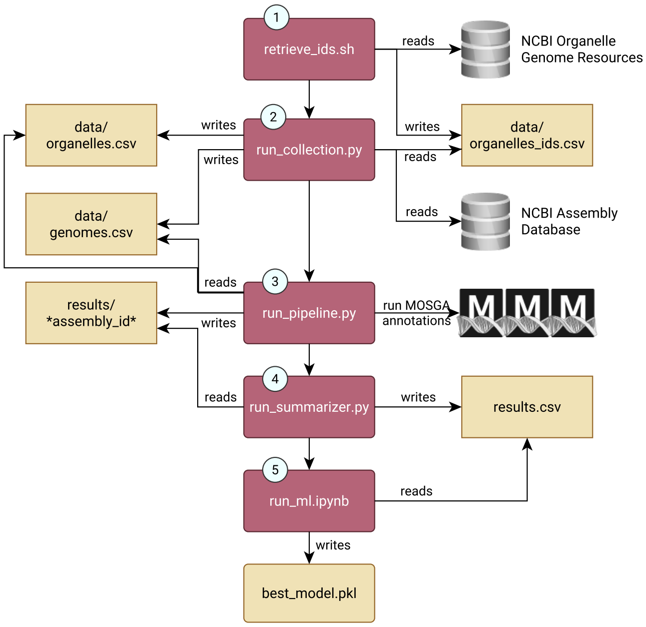


**Fig. S2. Data Retrieval and Machine Learning Workflow.** 1.) A bash script downloads all whole mitochondrial and plastid genomes from the NCBI Organelle Genome Resources Database. Extracts the FASTA sequence headers and write them into the data/organelles_ids.csv file. 2.) The identified sequence headers were searched in the NCBI Assembly database for corresponding genome assemblies containing organellar sequences in the summary description. 3.) All identified genome assemblies with a minimal and maximal threshold of included sequences were downloaded and annotated with one prepared MOSGA 2 configuration, including the relevant tools and parameterization. The organellar analysis summaries for each annotation run were stored, and corresponding organellar sequences were marked according to the previously identified organelles-to-assembly association. 4.) All results not failing during the annotation pipeline were summarized inside one results.csv data, which will be further used as the dataset for machine learning. 5.) Machine learning training and evaluation will be performed. The best model will finally be stored inside the best_model.pick file.

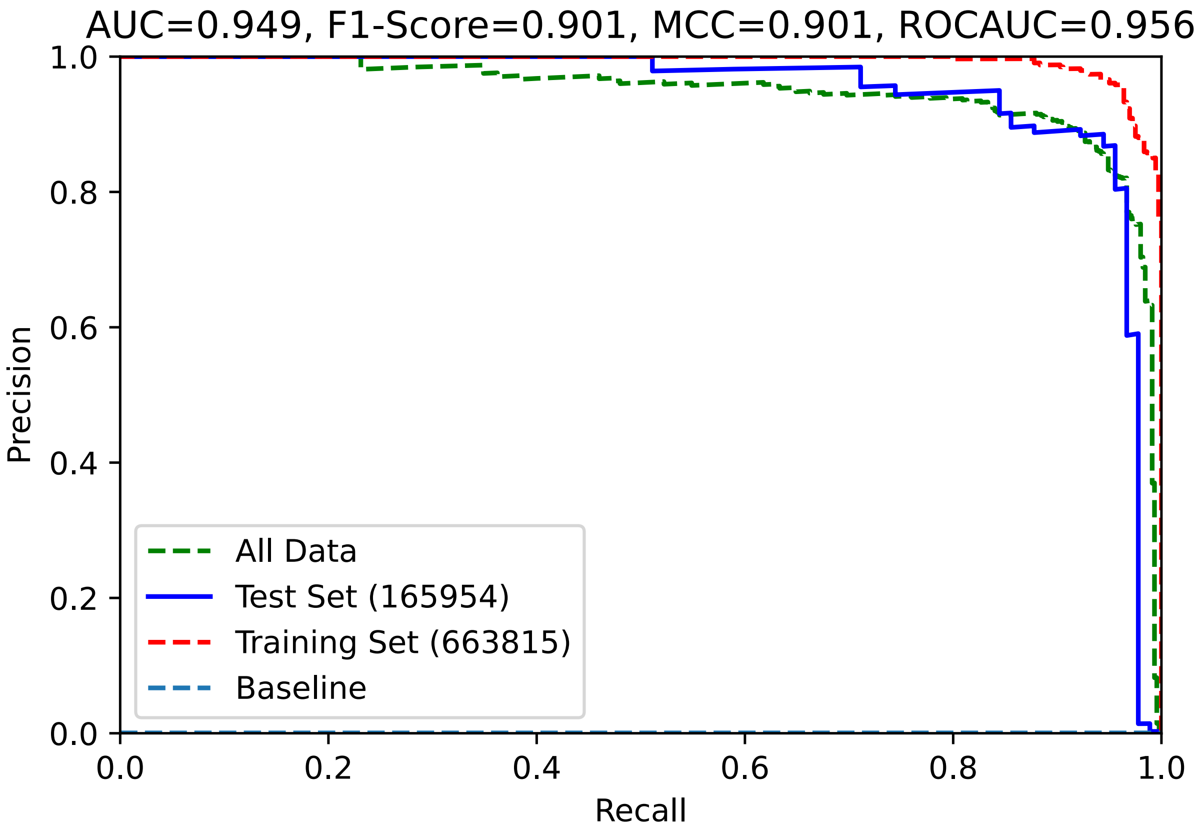


**Fig. S3. Precision-Recall curve.** This Precision-Recall curve of the finally selected model shows the tradeoff between the precision and recall on different thresholds.

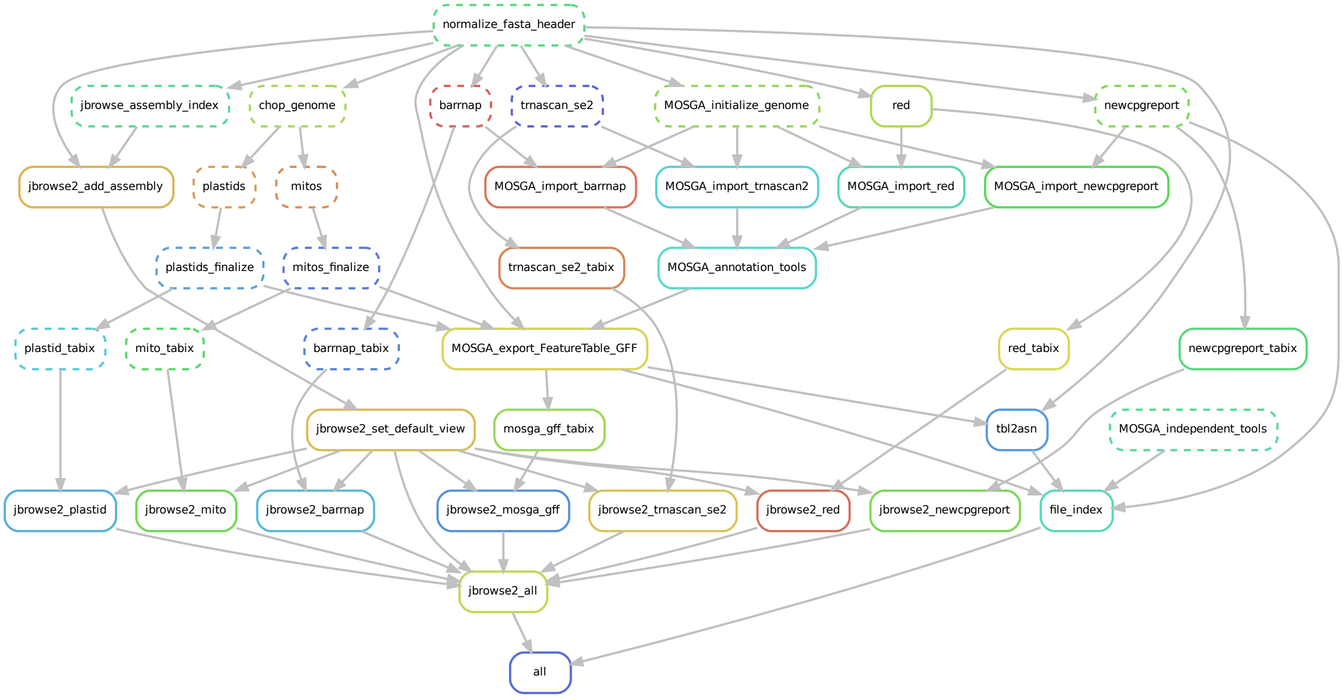


**Fig. S4. MOSGA 2 annotation workflow.** According to a prepared MOSGA 2 genome annotation workflow, MOSGA executed the visualized Snakemake rules to annotate required features that were used for machine learning.

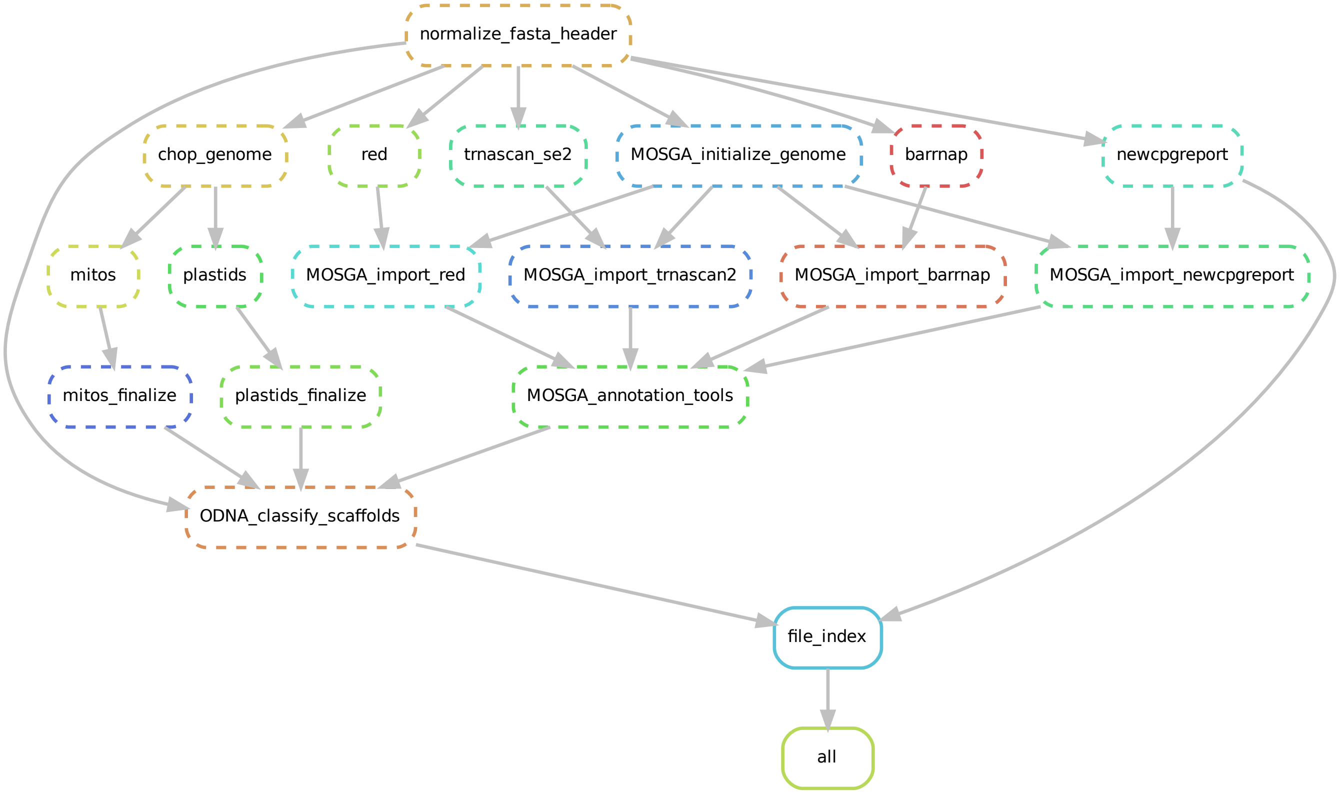


**Fig. S5. ODNA annotation workflow.** The minimalized genome annotation workflow that ODNA uses to classify each sequence within a genome.

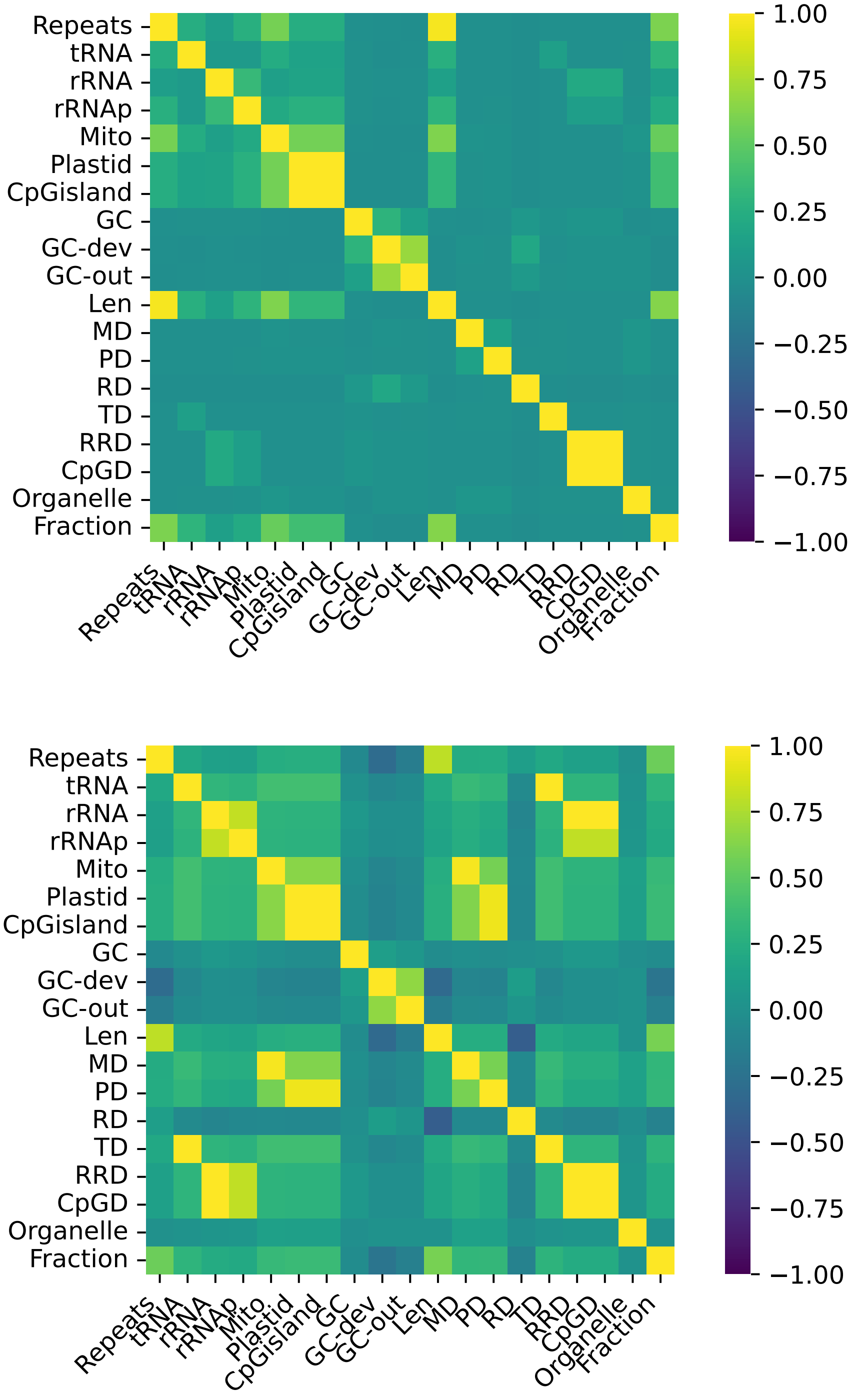


**Fig. S6. Feature correlation matrix.** The Pearson (a) and Spearman (b) correlation matrices show correlations between some features but no significant correlation between the target variable **Organelle** and any other feature.

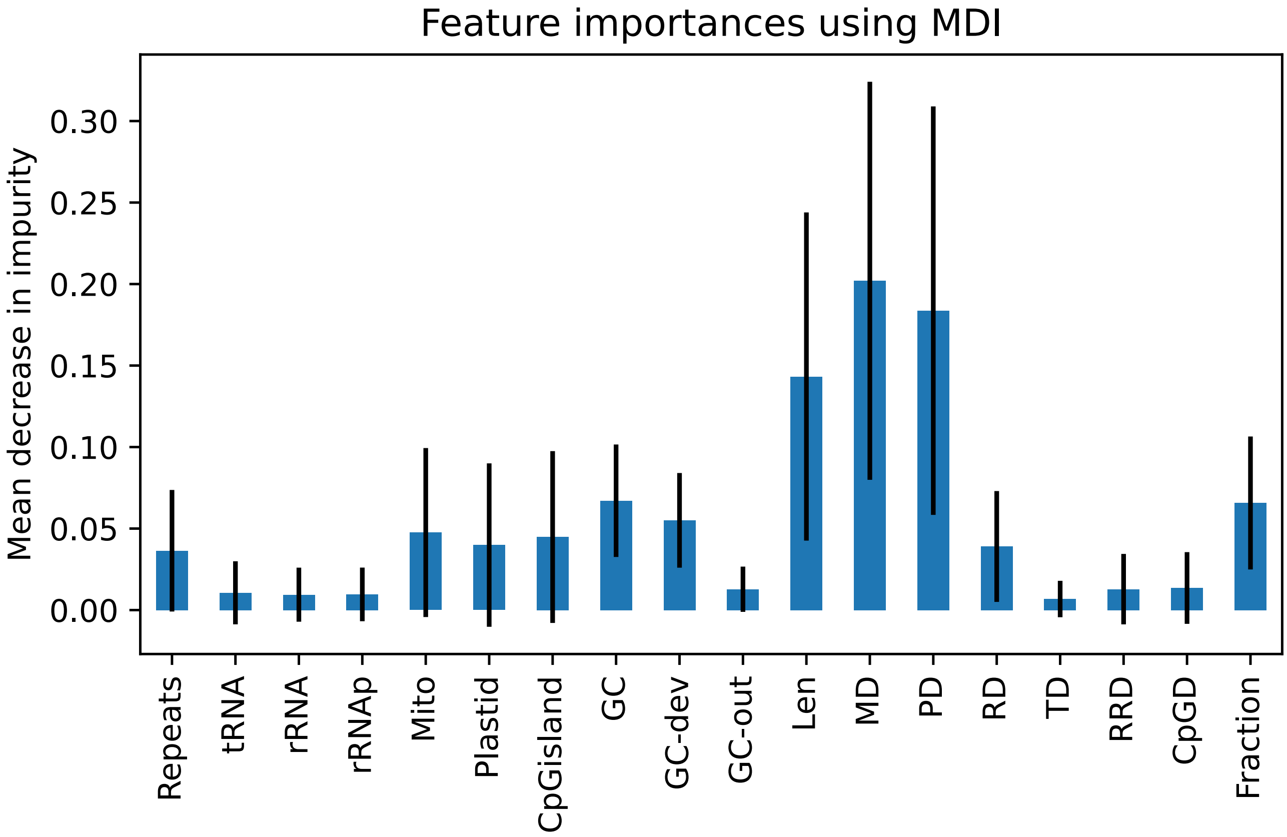


**Fig. S7. Mean decrease in impurity.** Although the final AdaBoost model was slightly superior to the random forest models, the mean decrease in impurity analysis from the random forest model reveals the most important features.

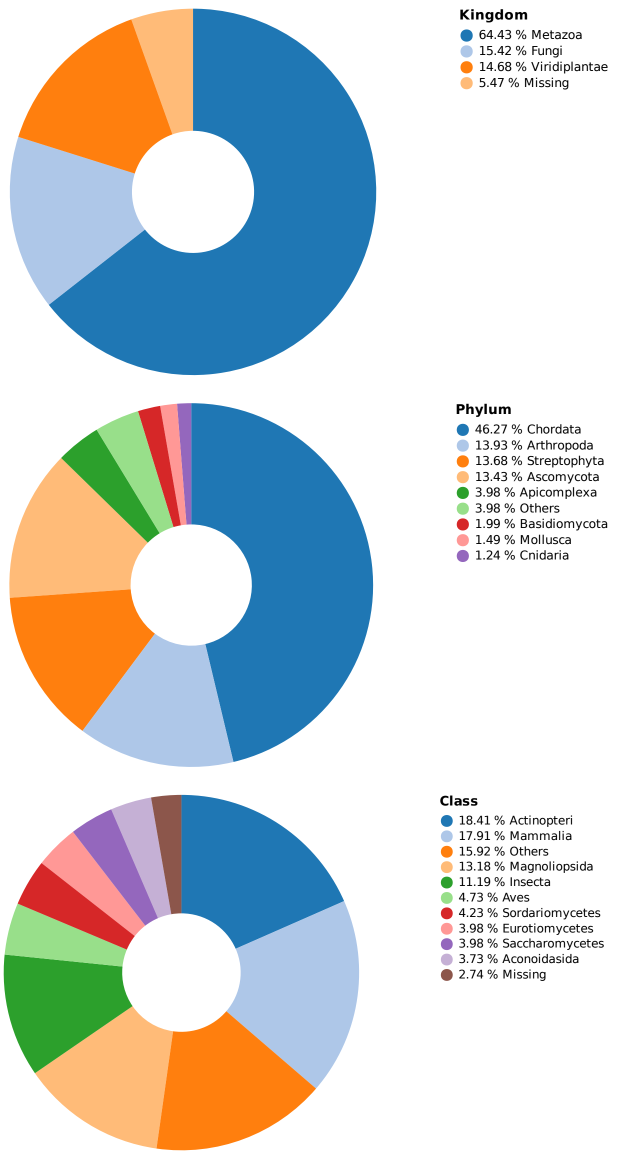


**Fig. S8. Taxonomic distribution of the used genomes categorized in the kingdoms (a), phyla (b), and classes (c).** Portions with less than two percent are summarized into the category "Others" while genomes with missing taxonomic information for a specific level are collected in the category "Missing". Species from the SAR clades, for example, do not have an assignment for a kingdom, phylum, or class.

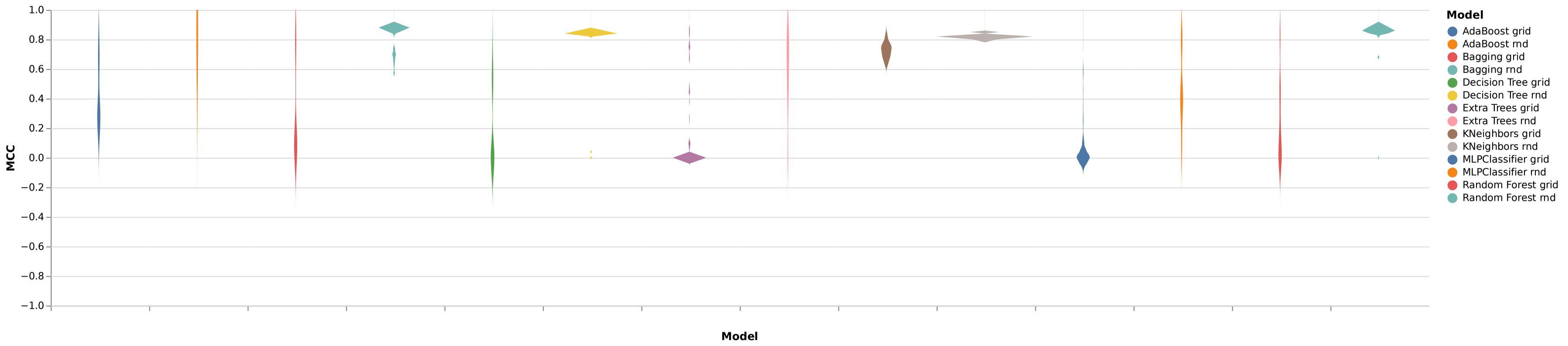


**Fig. S9. Hyperparameterization.** This violin plot shows the performance of all models generated with a 10-fold-cross-validation through hyper parameterization. The suffix in the model name indicates whether the HalvingGridSearchCV (*grid*) or RandomizedSearchCV (*rnd*) was used.

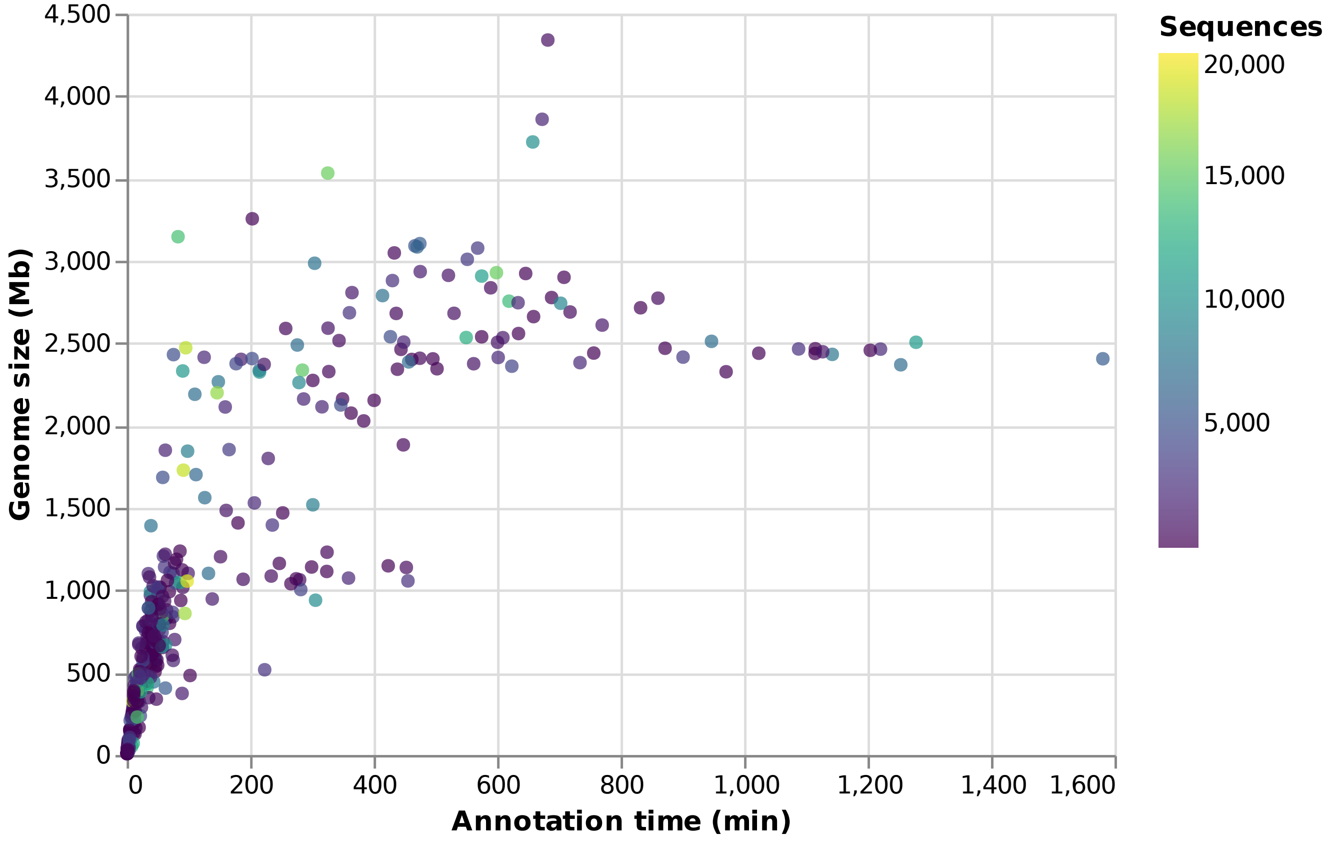


**Fig. S10. Genome annotation time in comparison to the genome size.** The total annotation time needed with two 32 cores servers roughly six weeks.
